# Transiently heritable fates and quorum sensing drive early IFN-I response dynamics

**DOI:** 10.1101/2022.09.11.507479

**Authors:** Laura C. Van Eyndhoven, Vincent P.G. Verberne, Carlijn V.C. Bouten, Abhyudai Singh, Jurjen Tel

## Abstract

Type I Interferon (IFN-I)-mediated antiviral responses are central to host defense against viral infections. Crucial is the tight and well-orchestrated control of cellular decision-making leading to the production of IFN-Is. Innovative single-cell approaches revealed that the initiation of IFN-I production is only limited to a fraction of 1-3% of the total population, both found *in vitro* and *in vivo*, which were thought to be stochastically regulated. To challenge this dogma, we addressed the influence of various host-intrinsic factors, both stochastic and epigenetic, on dictating early IFN-I responses. Hypomethylating drugs increased the percentage of responding cells. Next, with the classical Luria-Delbrück fluctuation test, we provided evidence that the fate of becoming a responding cell is transiently heritable. Finally, while studying varying cell-densities, we substantiated an important role for quorum sensing, which was verified by mathematical modeling. Together, this systems immunology approach opens up new avenues to progress the fundamental understanding of cellular decision-making during early IFN-I responses, which can be translated to other (immune) signaling systems, and ultimately will improve IFN-I based immune therapies.

## Introduction

Type I Interferon (IFN-I)-mediated responses are central to host defense against viral infections (Ivashkiv and Donlin, 2014; Mesev et al., 2019). Crucial is the tight and well-orchestrated control of cellular decision making leading to the production of IFN-I, as impaired response dynamics leads to the pathogenesis of a plethora of diseases that go beyond antiviral immunity only (Musella et al., 2017; Psarras et al., 2017; Zhang et al., 2020). Over the past decades, multilayered stochasticity driving cellular heterogeneity and subsequent cellular decision-making during IFN-I responses has become increasingly apparent (Rand et al., 2012; Van Eyndhoven et al., 2021b). In short, IFN-I responses are elicited by fractions of so-called first responding cells, also referred to as ‘precocious cells’ or ‘early responding cells’, which start the initial IFN-I production upon viral detection (Bauer et al., 2016; Hjorton et al., 2020; Patil et al., 2015; Shalek et al., 2014; Van Eyndhoven et al., 2021a; Wimmers et al., 2018). Their production is further enhanced via autocrine signaling, inducing a feedforward loop resulting in the upregulation of interferon regulatory factor (IRF) 7 and other signaling components (Honda et al., 2006). Besides, first responders elicit additional IFN-I production in so-called second responders, which are activated upon IFN-mediated paracrine signaling in combination with viral detection. These two major events have also been described as the early phase and later phase of IFN-I responses (Honda et al., 2006). Especially the regulation of the early phase is of increasing interest, because this phase is currently thought to orchestrate population-wide IFN-I signaling (Patil et al., 2015; Van Eyndhoven et al., 2021b).

Up till today it remains unclear whether cellular decision-making to become an IFN-I-producer during the early phase is as a stochastic process (e.g., dictated by host intrinsic factors, such as limiting levels of transcription factors and other signaling intermediates), or a deterministic process (e.g., dictated by epigenetics, leading to a predispositioning to perform certain cellular behaviors). The later phase seems mainly driven by stochastic processes, as the outcome is mainly dictated by host-intrinsic factors (e.g., intrinsic and extrinsic gene expression noise). These can be manipulated by overexpression of signaling intermediates, such as retinoic acid-inducible gene I (RIG-I), IRF3, and IRF7, leading to an increased overall production of IFN-Is (Harrison and Moseley, 2020; Zhao et al., 2012). In contrast, recent evidence suggests that the early phase could be dictated by determinism (Bagnall et al., 2020; Shaffer et al., 2020; Talemi and Höfer, 2018; Van Eyndhoven et al., 2021a). Accordingly, a transiently heritable gene expression program related to IFN-I signaling, including the expression of *DDX58 (RIG-I), IFIT1, PMAIP1*, and *OASL*, was discovered to be initiated only in fractions of unstimulated cells (Shaffer et al., 2020). In other words, it seems that among a population of cells, even before the occurrence of a viral infection, only a fraction of cells gets epigenetically programmed to become first responders. Accordingly, numerous studies characterized epigenetic control of IFN-I-related genes (e.g., *IFNβ)*, which may be of crucial importance during the early phase of IFN-I response dynamics (Daman and Josefowicz, 2021; Gao et al., 2021). In turn, the phenomenon of epigenetically programming of first responders could be prone to the effects of quorum sensing, allowing for adequate population-wide response dynamics accounting for differences in cell densities (Antonioli et al., 2019; Bardou et al., 2021; Doğaner et al., 2016; Muldoon et al., 2020; Schrom et al., 2020).

In this study, we addressed the influence of various host-intrinsic factors, both stochastic and epigenetic, in dictating early IFN-I responses. After having validated the fraction of first responders, which was remarkably similar to what has been observed and characterized in immune cells and other cell systems before, we assessed the three most important aspects of extrinsic and host-intrinsic stochasticity on cellular decision-making [i.e., varying viral loads, heterogeneous IRF7 levels, and fluctuations in cell-cycle states]. Using epigenetic drugs and the classical Luria-Delbrück fluctuation test, we challenged the dogma on stochasticity dictating early IFN-I responses, thereby proving heritability in responsiveness instead (Clark et al., 2021; Shaffer et al., 2020). Finally, we assessed the effects of quorum sensing driving population-wide responsiveness, which we substantiated with an ordinary differential equation (ODE) model. Together, this systems immunology approach highlights the ability to challenge the fundamentals of cellular decision-making during early IFN-I responses, and potentially other immune signaling systems. Ultimately, these novel insights pave the way towards improved IFN-mediated immune therapies.

## Results

### Reporter cell model to study early IFN-I responses

Studying IFN-I dynamics in (human) primary immune cells allows for translation towards clinical applications, however, experimental approaches are often limited by relatively low cell counts, possible immune cell impurities, and additional layers of stochasticity introduced by the presence of heterogeneous subsets (Van Eyndhoven et al., 2021a). Therefore, we utilized murine reporter cells to provide us with a robust model to study early IFN-I responsiveness (Rand et al., 2012). The early IFN-I phase is characterized by the detection of viral nucleic acids by pathogen recognition receptors, leading to the phosphorylation and translocation of IRFs (e.g., IRF3 and IRF7) from the cytoplasm to the nucleus, where they initiate the transcription of IFN-Is (Honda et al., 2006; Rehwinkel and Gack, 2020) (Fig. 1a). Subsequently, the later phase is characterized by the signaling induced by IFN-Is activating IFN-I receptors (IFNARs). This leads to the phosphorylation, complex formation, and translocation of signal transducer and activator of transcription 1 (STAT1), STAT2, and IRF9, termed IFN-stimulated gene factor 3 (ISGF3), to initiate the transcription of interferon stimulated genes (ISGs). Accordingly, we used a NIH3T3-IRF7-CFP reporter cell line, which constitutively expresses fluorescent IRF7 molecules, to monitor signaling dynamics during the early phase of the response (Fig. 1b). For this cell model, IRF7 translocation correlates with IRF3 translocation, making IRF7 translocation as only readout sufficient to study first responders (Rand et al., 2012). The NIH3T3-STAT1-CFP/STAT2-YFP reporter cell line was utilized later to validate the production of IFN-Is upon translocation of IRF7.

**Figure 1.**
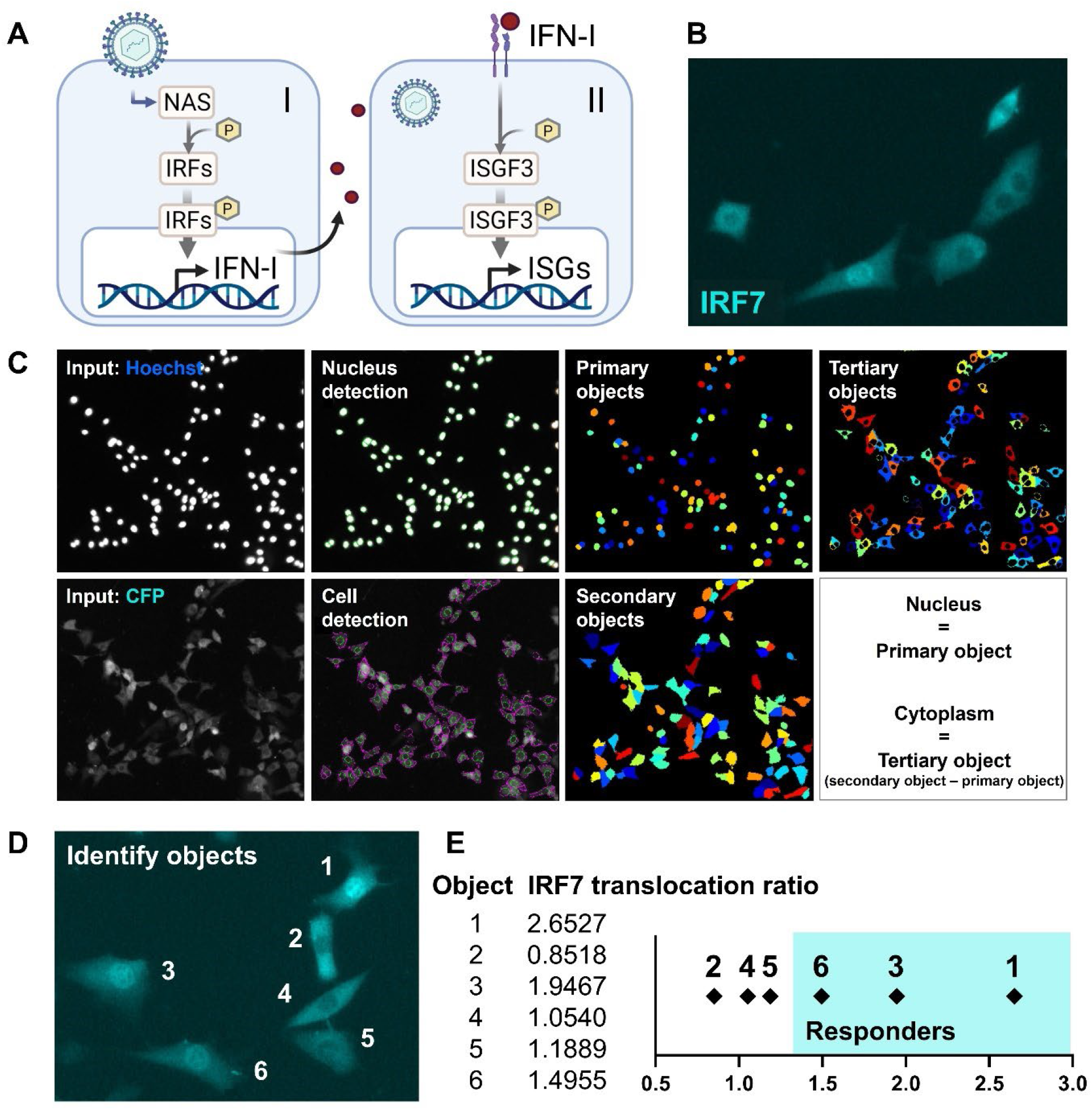
Reporter cell model to study early IFN-I responses. **A** Schematic overview of the early (I) and later (II) phase IFN-I responses. The early phase is characterized by the detection of nucleic acids by nucleic acid receptors (NAS), followed by the phosphorylation (p) and translocation of interferon regulatory factors (IRFs) and subsequent induction of IFN-Is. Upon paracrine signaling, IFN-Is bind to IFN-I receptors (IFNARs), leading to the phosphorylation and translocation of interferon stimulated gene factor 3 (ISGF3), consisting of STAT1, STAT2, and IRF9, respectively, inducing the production of interferon simulated genes (ISGs). **B** Microscopy image of NIH3T3 cells stably expressing the fusion protein IRF7–CFP. **C** Image processing and analysis steps in CellProfiler script for the detection of fluorescent signal in the nuclei and cytoplasms. **D** Example image with 6 identified objects. **E** IRF7 transfection ratios of example objects plotted.

To identify first responders, translocation of fluorescently tagged IRF7 was monitored in an unbiased fashion using a custom-made automated image analysis script developed in CellProfiler software (Fig. 1c, Supplementary Fig. 1a-c) (Stirling et al., 2021). Primary objects (nuclei) were detected and defined based on the Hoechst signal after nuclei staining. Next, the secondary objects (cells) were detected and defined based on the CFP signal originating from the fluorescently tagged IRF7 molecules. Finally, the tertiary objects (cytoplasms) were defined by subtracting the primary objects from the secondary objects.

First responders were defined by determining the IRF7 translocation ratio by dividing the CFP median intensity from the cytoplasm by the CFP median intensity from the nucleus (N/C) (see Materials and Methods). As an example, 6 cells were imaged simultaneously, containing three responding cells showing clear translocation of signal, and three non-responding cells showing relatively less signal inside the nucleus (Fig. 1d). Indeed, the three cells that would have been defined by eye as responding cells had the highest IRF7 translocation ratio (Fig. 1e, Supplementary Fig. 2a-d).

Together, we established the detection of first responding cells in a high-throughput, unbiased manner, based on the translocation of fluorescent signal corresponding with IRF7 molecules from the cytoplasm to the nucleus.

### Validation of first responders in a reporter cell model

To elicit early IFN-I responses in our model, we used rhodamine-labeled Poly(I:C), instead of live or attenuated viruses, thereby avoiding any additional stochasticity introduced by viral extrinsic factors (e.g., genetic variability among the virus population, variability in viral replication, etc.). By using rhodamine-labeled Poly(I:C) over regular Poly(I:C), we were able to carefully track transfection efficiencies over time (Fig. 2a, b). To limit noise introduced to the system, resulting from poor transfection timing and efficiencies, we optimized transfection to achieve fast and potent delivery of stimulus (Fig. 2c, Supplementary Fig. 3).

**Figure 2.**
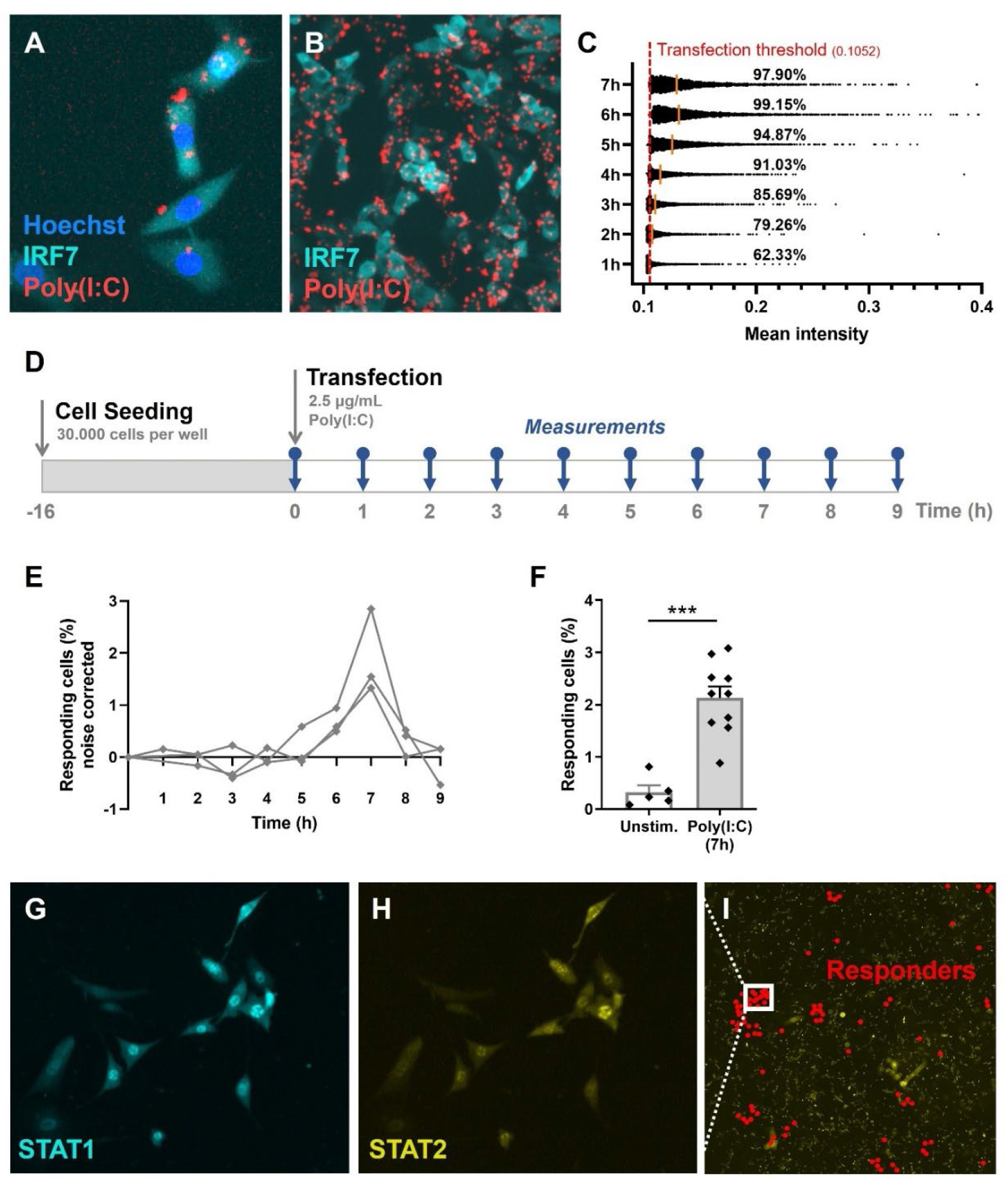
Validation of first responders in reporter cell model. **A** Microscopy picture of NIH3T3: IRF7-CFP, stained with Hoechst nuclear stain, transfected with rhodamine-labeled Poly(I:C). **B** Overview of transfected cells. **C** Transfection efficiency quantification over time, based on rhodamine mean intensity detected in cells. **D** Experimental design of first responder validation in NIH3T3: IRF7-CFP cells. **E** Percentages of noise corrected responding cells. Cells were seeded on coverslips 16 hours prior to transfection with 2.5 μg/mL Poly(I:C). Over the first 9 hours, the percentages of translocated cells were determined (n = 3 experimental replicates). **F** Percentages of responding cells after 7 hours of Poly(I:C) transfection, compared to unstimulated cells (n = 10; p = 0.0003). **G** Microscopy image of NIH3T3: STAT1-CFP; STAT2-YFP, for additional first responder validation. Cells were seeded and transfected as described before. Translocation of STAT1 was assessed after 7 hours post transfection. **H** Corresponding image of STAT2-YFP signal. **I** Corresponding overview image of population of NIH3T3: STAT1-CFP; STAT2-YFP, with responding (translocated) cells indicated with red dots. Data information: in (F), data are represented as mean ± SEM. ***P≤0.001 (Mann-Whitney test).

Next, we set out to explore the response dynamics over the first 9 hours post transfection to determine the response peak (Fig. 2d). Earlier studies indicated a peak of IRF7 translocation around 8 hours, and a peak of IFN-beta (IFNβ) production around 10 hours post activation [i.e., using Poly(I:C) and Newcastle Disease Virus] (Rand et al., 2012). Accordingly, upon transfection optimization, in our experiments the response peaked at 7 hours post transfection, with an average of 2.1% of responding cells (Fig. 2e, f). This percentage is in line with what has been found across literature, species [i.e., human and mice] and cell types [i.e., fibroblasts, monocyte derived dendritic cells, plasmacytoid dendritic cells], emphasizing the elegant yet robust feature of only a fraction of first responding cells driving the population-wide IFN-I system (Bauer et al., 2016; Drayman et al., 2019; Patil et al., 2015; Shalek et al., 2014; Van Eyndhoven et al., 2021a; Wimmers et al., 2018). Besides, the background numbers of translocated cells possibly reflect the intrinsic feature of the IFN-I system to ensure basal IFN-I expression and IFNAR signaling to equip immune cells to rapidly mobilize effective antiviral immune responses (Ivashkiv and Donlin, 2014).

Accordingly, we wondered whether we could capture the orchestrating role of first responders on population-wide IFN-I response dynamics. Therefore, we studied the response dynamics using a NIH3T3-STAT1-CFP/STAT2-YFP fibroblast reporter cell line. Seven hours post infection, IFNβ produced by the first responders will diffuse to neighboring, yet nonresponding cells, thereby activating their IFNARs, followed by the subsequent translocation of ISGF3, consisting of STAT1, STAT2 and IRF9. Accordingly, at 7 hours post infection, we were able to capture clusters of STAT1/STAT2 translocated cells (Fig. 2g, h, i). These clusters represent a phenomenon of competition between cytokine diffusion [i.e., IFN-Is produced by first responder] and consumption [i.e., by surrounding cells] generating spatial niches of high cytokine concentrations with sharp boundaries (Oyler-Yaniv et al., 2017).

Taken together, we established a methodology for rapid and potent delivery of stimulus, thereby minimizing the potential noise introduced by extrinsic factors, to further reveal the multilayered stochasticity driving first responders. Additionally, we validated the presence of fractions of first responders, and validated their ability to induce population-wide IFN-I signaling.

### Extrinsic and intrinsic stochasticity dictating first responders

In contrast to the role of host-intrinsic factors, literature stated that the role of extrinsic factors (those that are introduced by the virus/stimulus itself) is rather small in determining the fraction of first responders, indicated by the lack of dose-dependent effects and the robustness of percentages of first responders across stimulus types (Shalek et al., 2014; Van Eyndhoven et al., 2021a; Wimmers et al., 2018). Of note, on the contrary, extrinsic factors can correlate with the percentage of second responders, though studies often do not distinguish between these two different cell fates, but focusing on population-wide responses instead (Rand et al., 2012; Zhao et al., 2012). To test the effect of a variety of extrinsic factors on the first responders, we first tested for a correlation between the responsiveness [i.e., IRF7 translocation ratio] and the actual amount of stimulus received by the cells, which was only very low, though significant (R^2^ = 0.0171, p < 0.0001) (Fig. 3a). Of note, the responsiveness does not reflect the fraction of responders, which is hypothesized to be unaffected by stimulus dosage, as observed in plasmacytoid dendritic cells upon single-cell stimulation in microfluidic droplets (Wimmers et al., 2018). Therefore, we conclude that first responders, especially the fractions, are only minorly influenced by stimulus dosage.

**Figure 3.**
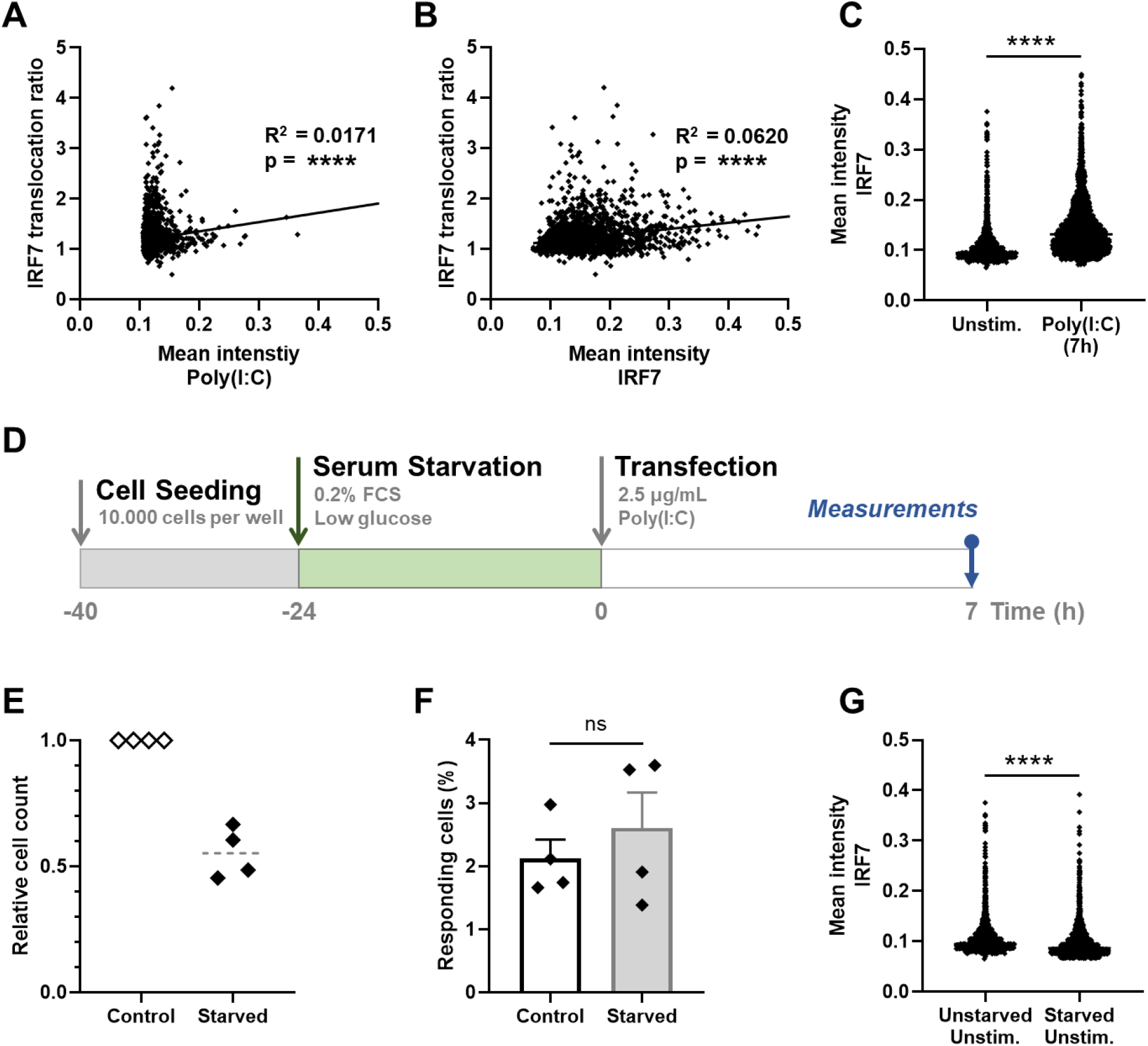
Extrinsic and intrinsic stochasticity dictating early IFN-I responses. **A** NIH3T3: IRF7-CFP cells were seeded and transfected as described before. At 7h post transfection, images were analyzed using an automated image analysis script to measure rhodamine-labeled Poly(I:C) intensities, and the IRF7 translocation ratios. Plotted are the mean intensities of Poly(I:C) against the IRF7 translocation ratios (R_2_ = 0.0171). **B** As in panel (**a**), the mean intensities of total IRF7 were measured. Plotted are the background levels of IRF7 against the IRF7 translocation ratios (R_2_ = 0.0620). **C** Scatter plot depicting the IRF7 levels of unstimulated cells versus Poly(I:C) stimulated cells after 7 hours (p < 0.0001). **D** Experimental design of serum starvation experiments in NIH3T3: IRF7-CFP cells. Cells were seeded 40 hours prior to the start of the experiment. 24 hours prior to transfection, cells were serum and glucose deprived. Next, cells were transfected with 2.5 μg/mL Poly(I:C) and assessed for nuclear translocation of IRF7 after 7 hours. **E** Validation of cell cycle arrest induced by serum starvation by relative cell counts of the control (unstarved) conditions, compared to the corresponding starved conditions (n = 4). **F** Comparison of the percentages of responding cells of the control conditions, compared to the starved conditions (nonsignificant = ns). **G** Scatter plot of a representative biological replicate comparing the IRF7 levels of unstimulated cells, as in (**c**), versus starved, unstimulated conditions (p < 0.0001; Mann-Whitney test, two-tailed). Data information: in (F), data are represented as mean ± SEM. ****P≤0.0001 (Mann-Whitney test).

Like all biochemical reactions, stochastic processes (e.g., gene expression noise) influence IFN-I response dynamics. Universally, intrinsic gene expression noise results from the stochastic nature of biochemical reactions, whereas extrinsic gene expression noise results from cell–cell fluctuations of components that are involved in generating the response (Dey et al., 2015). In essence, every step of IFN-I signaling involves limiting signaling intermediates, making every step subject to the effects of gene expression noise (Zhao et al., 2012). While IRF7 is one of the key factors driving IFNβ production, and thereby possibly driving the first responders, we set out to investigate the relation between background levels of IRF7 and first responding cells. However, we could only find a very weak correlation between IRF7 translocation ratio and IRF7 mean intensity (R^2^ = 0.0620, p < 0.0001), arguing that first responders are only minorly driven by differences in background levels of IRF7 (Fig. 3b). Although, it was interesting to observe the degree of heterogeneity in background IRF7 expression levels, as well as the significant increase in signal after stimulation (Fig. 3c). The latter confirms the already well characterized feedback loops enhancing the IRF7 expression after autocrine and paracrine signaling induced by the first responders.

Next, we wondered whether cell cycle state could be a potential driver, since studies pointed towards a role for cell cycle state dictating IFN-I production, though mainly related to second responding cells (Cingöz and Goff, 2018; Mudla et al., 2020). To elucidate the effect of cell cycle state on first responding cells, we synchronized the cells using serum starvation for 24 hours (Fig. 3d). This approach induces a cell cycle arrest, halting cells in the G0/G1 phase, thereby synchronizing the whole population (Chen et al., 2012). We validated the cell cycle arrest by comparing the cell counts of starved conditions with unstarved conditions, which in theory should differ a factor of 2, knowing the cells divide approximately every 24 hours. Indeed, only half of the cell numbers could be detected after 24-hour serum starvation, compared to the corresponding control samples (Fig. 3e). Interestingly, the percentage of first responding cells obtained from the starved conditions did not significantly differ from the percentages obtained from the unstarved conditions, suggesting that there is no significant effect of cell cycle state on first responders (Fig. 3f). Additionally, the background levels of IRF7 were significantly lower for the starved conditions, compared to the unstarved conditions, again validating our successful approach of starving the cells, which limits the overall protein synthesis (Fig. 3g).

In short, the extrinsic and intrinsic factors that were assessed in this study turned out to be only minorly dictating the cellular decision to become a first responder. Of note, these results do not exclude other (extrinsic or intrinsic) factors (e.g., those involved in the phosphorylation and translocation of IRF7), those that were not included in this study, from playing important roles in dictating first responders.

### Epigenetic regulation dictating first responders

Our results thus far proved that stochastic features are only minorly driving first responders, which made us further explore the influence of deterministic features instead. Importantly, although the terms stochasticity and determinism seem highly dichotomous, deterministic features (e.g., epigenetic regulation) are often, if not always, stochastically regulated (Zernicka-Goetz and Huang, 2010). However, in cellular decision-making, the major difference between a stochastic process and a deterministic process boils down to the effects of (varying) inputs on dictating (varying) outputs. In fact, a stochastic process in characterized by the exact same stimulus leading to varying response outcomes, often as a result of varying host-intrinsic factors (Symmons and Raj, 2016). In contrast, a deterministic process is characterized by an outcome (e.g., IFN-I production) that is fixed, or at least to a large degree, while the input can be variable. How cells are epigenetically predispositioned, in turn, can again be a stochastic process, similar to the fundamentals of developmental biology in which cells are randomly pushed towards deterministic outcomes (Zernicka-Goetz and Huang, 2010).

Remarkably, throughout the experiments we observed the occurrence of two neighboring cells showing translocation (Fig. 4a, b, Supplementary Fig 4a-d). If being a first responder is stochastically regulated, the probability of one of the neighboring cells also being a responder is remarkably small, knowing the response rate is only 2.134%. In fact, assuming a cell has on average 4 neighboring cells, the probability of at least one of them being a responder equals the probability of 1−1−*none responds*=1 − 0.9786^4^=0.0826=8.26%. Therefore, the observation of so-called responding sister cells further supported the hypothesis that first responders are predetermined. In other words, it seemed more likely that cells that were predispositioned to become a first responder passed this on to their daughter cells, that upon activation both show translocation. Also, after realizing that in the general experimental setup cells were seeded approximately 24 hours before imaging, allowing all cells to have divided once by the time of imaging, the appearance of responding sister cells could be further explained and quantified. Interestingly, comparing the two responding sister cells, the background levels of IRF7 can differ drastically (Fig. 4a), but can also be remarkably similar (Fig. 4b).

**Figure 4.**
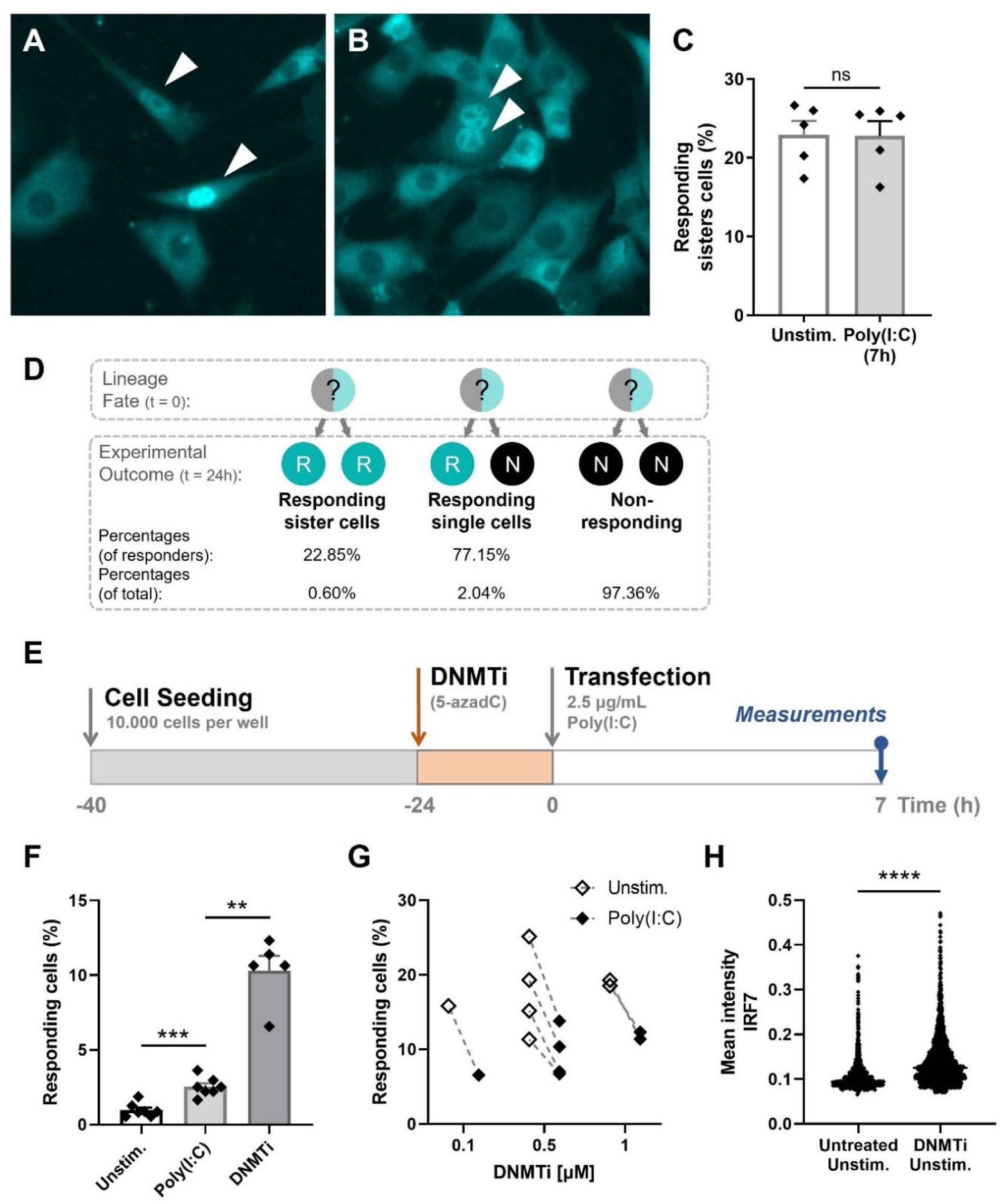
Epigenetic regulation dictating early IFN-I responses. **A** NIH3T3: IRF7-CFP cells were seeded on coverslips and transfected with 2.5 μg/mL Poly(I:C) for 7 hours. Microscopy image of two responding, neighboring cells, referred to as responding sister cells, displaying different background levels of IRF7. **B** Microscopy image of two responding sister cells, displaying similar background levels of IRF7. **C** Data on percentages of responding sister cells for unstimulated conditions (background translocation) versus stimulated conditions, transfected with Poly(I:C) after 7 hours. **D** Schematic of theoretical lineage fates and subsequent experimental outcomes (depicted as percentages of responders and of total population) upon cellular division. **E** Experimental design of epigenetics experiments in NIH3T3: IRF7-CFP cells. Cells were seeded 40 hours prior to the start of the experiment. 24 hours post transfection, cells were treated with DNMTi to induce hypomethylation. Next, cells were transfected with 2.5 μg/mL Poly(I:C) and assessed for nuclear translocation of IRF7 after 7 hours. **F** Percentages of responding cells for unstimulated, stimulated (Poly(I:C)), and DNMTi (1 μM) treated + stimulated conditions. **G** Data on paired percentages of responding cells (unstimulated versus stimulated) for different concentrations of DNMTi. **H** Scatter plot of a representative biological replicate comparing the IRF7 mean intensity of individual cells of untreated, unstimulated conditions, versus DNMTi treated, unstimulated conditions. **P≤0.01, ***P≤0.001, ****P≤0.0001. Data information: in (C,F), data are represented as mean ± SEM. **P≤0.01, ***P≤0.001, ****P≤0.0001 (Mann-Whitney test).

Next, we quantified the percentage of responding sister cells (neighboring cells) for both the unstimulated (observed background translocation levels) and stimulated conditions, which were not significantly different from one another (Fig. 4c). The criteria for responders being assigned as responding sister cells included a maximum distance between the two cells of 300 μm, and a maximum of one nonresponding cells between the two responders. In theory, with an average of 22.85% of responding sister cells, it implies that two responding sister cells originated from one mother cell in 22.85% of the cases (Fig. 4d). In 77.15% of the cases, only one of the two sister-cells turned out to become a first responder. For this scenario, it is yet unclear whether the potential transfer of responder fate (assuming the mother cell was a responder) was only succeeded for only one daughter cell, or whether this single responding daughter appeared stochastically from a nonresponding lineage (assuming the mother cell was a nonresponder). Both have been described in literature, referred to as transiently heritable cell fates (Shaffer et al., 2020).

Continuing the hypotheses of transiently heritable cell fates stated in literature, we investigated the manipulation of cellular decision-making by altering the cells’ epigenetic profile, thereby altering any potential predispositioning towards becoming a first responder. Therefore, cells were incubated with DNA methyltransferase inhibitor (DNMTi) 5-Aza-2′-deoxycytidine (5-azadC) 24 hours prior transfection (Fig. 4e). We hypothesized that, under regular circumstances, in only 1-3% of cells the epigenetic profiling allows the cell to become a first responder. Accordingly, hypomethylating the DNA of all cells consequently will result in higher response rates. Indeed, cells treated with DNMTis showed higher percentages of first responding cells, arguing that the cellular decision to become a responder is, at least partly, epigenetically regulated via DNA methylation (Fig. 4f). However, unstimulated cells treated with DNMTis also showed increased percentages of first responders, with even higher percentages compared to the stimulated DNMTi-treated cells (Fig. 4g). This might be explained by the effect of DNMTis triggering cytosolic sensing of double stranded RNA originating from retroviruses, that are no longer silenced while using these types of drugs (Chiappinelli et al., 2015). We later confirmed this by showing increased levels of IRF7 mean intensities in unstimulated, DNMTi-treated cells, compared to unstimulated untreated cells (Fig. 4h). Namely, this implies that, though these cells were not transfected with Poly(I:C), these cells got properly activated by the retroviruses, leading to the subsequent production of IFNβ, thereby initiating the positive feedback loops causing higher IRF7 expression levels. A plausible explanation for the decreased response numbers for the Poly(I:C)-activated cells compared to the unstimulated cells is that a more potent nucleic acid signaling (the combination of Poly(I:C) and retroviruses) elicits a stronger negative feedback, compared to the activation induced upon retroviruses only, either directly or indirectly via the upregulated ISG expression upon genome-wide DNA hypomethylation.

Taken together, we show that, at least partly, the cellular decision-making to become a first responder is epigenetically regulated via methylation. Although the self-activation by retroviruses might be considered as an artifact, the results still prove that upon hypomethylation and activation [i.e., either by only retroviruses or in combination with Poly(I:C)], the fraction of first responders increases.

### Fluctuation analysis on first responders

Another elegant approach to assess whether epigenetic mechanisms are involved in driving first responders involves the classical Luria-Delbrück fluctuation test (Luria and Delbrück, 1943). It was originally used to demonstrate the occurrence of genetic mutations in bacteria in the absence of selection, rather than being a response to selection, in which variability between different clonal populations is assessed. Similarly, a stochastic feature would be equally present among different clones, whereas a (transiently) heritable feature can widely fluctuate between different clones, depending on the cell fate of the mother cells (Shaffer et al., 2020).

Assuming first responders are purely stochastically regulated, probability calculations can predict from which generation number the probability of at least one first responder present is close to one, knowing that on average only 2.134% first responders are present in a population (see Materials and Methods). From generation 6 onwards, the probability of at least one responder being present becomes considerably high (Fig. 5a). Subsequently, each clone, consisting of ∼ 64 cells, would have 1.37 responding cells on average (Fig. 5b). Therefore, we considered clones of generation 6 and older to be most informative, taking into account the trade-off between risking the absence of first responders because of cell numbers being simply too low, and risking clones that again start representing regular cultures (cells that have been kept in culture for multiple passages according to conventional cell culture). For the latter, we decided to compare clones of generation 6 with clones of generation 9, 13, 16, and with results obtained from regular cultures (generation ∞). Of note, in this experimental setting, the generation number is only an indication of the amount of cellular divisions that the clone has undertaken, rather than a determinantal factor, as cells do not remain synchronized over multiple generations.

**Figure 5.**
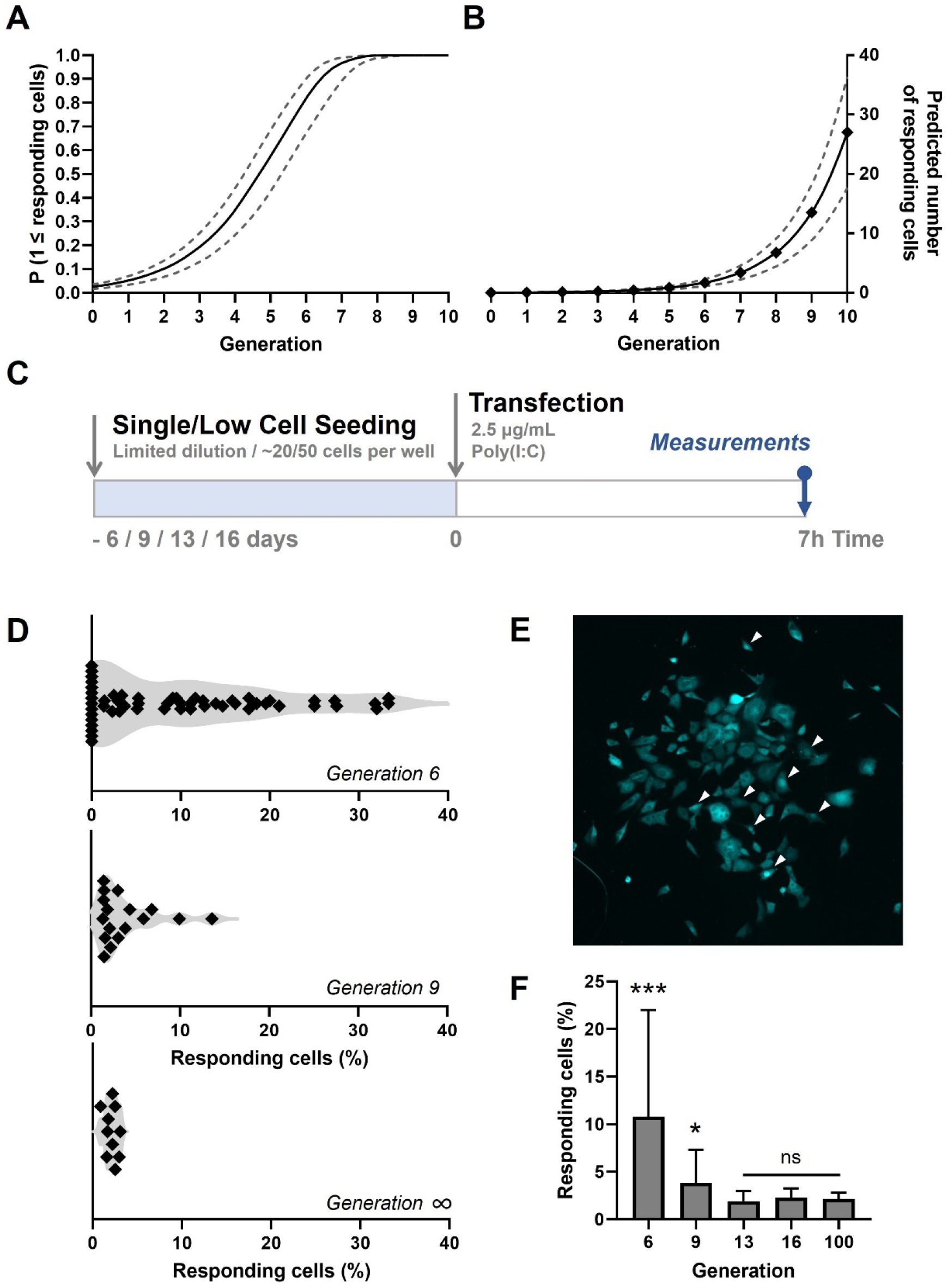
Fluctuation analysis on early IFN-I responses. **A** Probability curves for the presence of at least one first responder per clone over the first 10 generations, assuming stochasticity. Solid line represents the probability based on the mean percentage of first responders in regular cultures; dashed lines represent the mean plus and minus the standard deviation (SD). **B** Curves on predicted numbers of first responders present over the first 10 generations, based on the mean percentages obtained from regular cultures (solid line), and the mean +/-SD (dashed line). **C** Experimental design of fluctuation experiments in NIH3T3: IRF7-CFP cells. Cells were either seeded following limited dilution 20 hours prior to the start of the experiment, or at only ∼50 cells per 24-well 6 days prior to the start of the experiment. Next, cells were transfected with 2.5 μg/mL Poly(I:C) and assessed for nuclear translocation of IRF7 after 7 hours. **D** Fluctuation plots on percentages of responding cell of clones of generation 6 (n = 56 clones), 9 (n = 17 clones) and *∞* (regular cultures; n = 10). **E** Microscopy image of clone of 6 generations displaying numerous translocated cells, some of which are indicated with white arrows. **F** Percentages of responding cells of clones of different generations. Data information: in (F), data are represented as mean ± SD; Welch’s t test, two-tailed; ****p < 0.0001 ; *p = 0.0446.

To generate the clones of generation 13 and 16, we used conventional limited dilution approaches to seed single cells in single micro-wells (Fig. 5c). The clones of 6 and 9 generations were generated by seeding only approximately 20 or 50 cells per 24-wells on coverslips, respectively, to assure optimal culture conditions during the very first and critical days of cloning. In practice, after 6 or 9 days, there is still enough empty space surrounding the clusters of cells to determine which cells originated from a single cell (Supplementary Fig. 5a-c). Next, clones were stimulated and checked for first responders as described before. Remarkably, some clones of 6 generations showed over 20% of first responding cells, whereas other clones showed no single translocation event (Fig. 5d, e, Supplementary Fig. 5a-d). In other words, we observed a rather large variation in percentages of responding cells in clones of generation 6. Besides, the mean across clones from generation 6 was significantly higher than observed in regular cultures, which convincingly proves that first responders are not purely stochastically regulated, as this would universally lead to similar fractions throughout (Fig. 5f). Interestingly, clones of generation 9 still showed a fluctuation which was higher than what was observed in regular cultures, but it was already remarkably less than clones of generation 6, with no clones showing 0 responders (Supplementary Fig. 6a-d). From clones of generation 13 onwards, percentages were not significantly different from regular clones.

These results further confirm that first responders are epigenetically regulated and that there is a rather high heritable factor involved.

### Modeling cellular-decision making during early IFN-I responses

For a proper interpretation of the results obtained from the fluctuation assay, we modeled cellular decision-making during early IFN-I responses, where individual cells are either displaying IRF7 translocation, making them first responders (ON), or not (OFF). Assuming a purely stochastic process, upon cloning, the total mean across clones should be equal to the mean obtained from regular cultures, which would be 2.134% (Fig. 6a). Accordingly, the coefficient of variation (CV) is determined by the biological and technological variation, therefore considered relatively low. The rate in which responders appear in the population (k_on_) is also relatively low, corresponding with the probability of a cell to become a responder (p = 0.02134). Assuming a strictly heritable fate, meaning that all responding cell will divide into responding daughter cells, the total mean across clones will not change (Fig. 6b). However, the CV will be much higher than the biological and technological noise, determined by the occurrence of responding lineages. The kon is not defined, as individual cells will no longer change fate across the generations.

**Figure 6.**
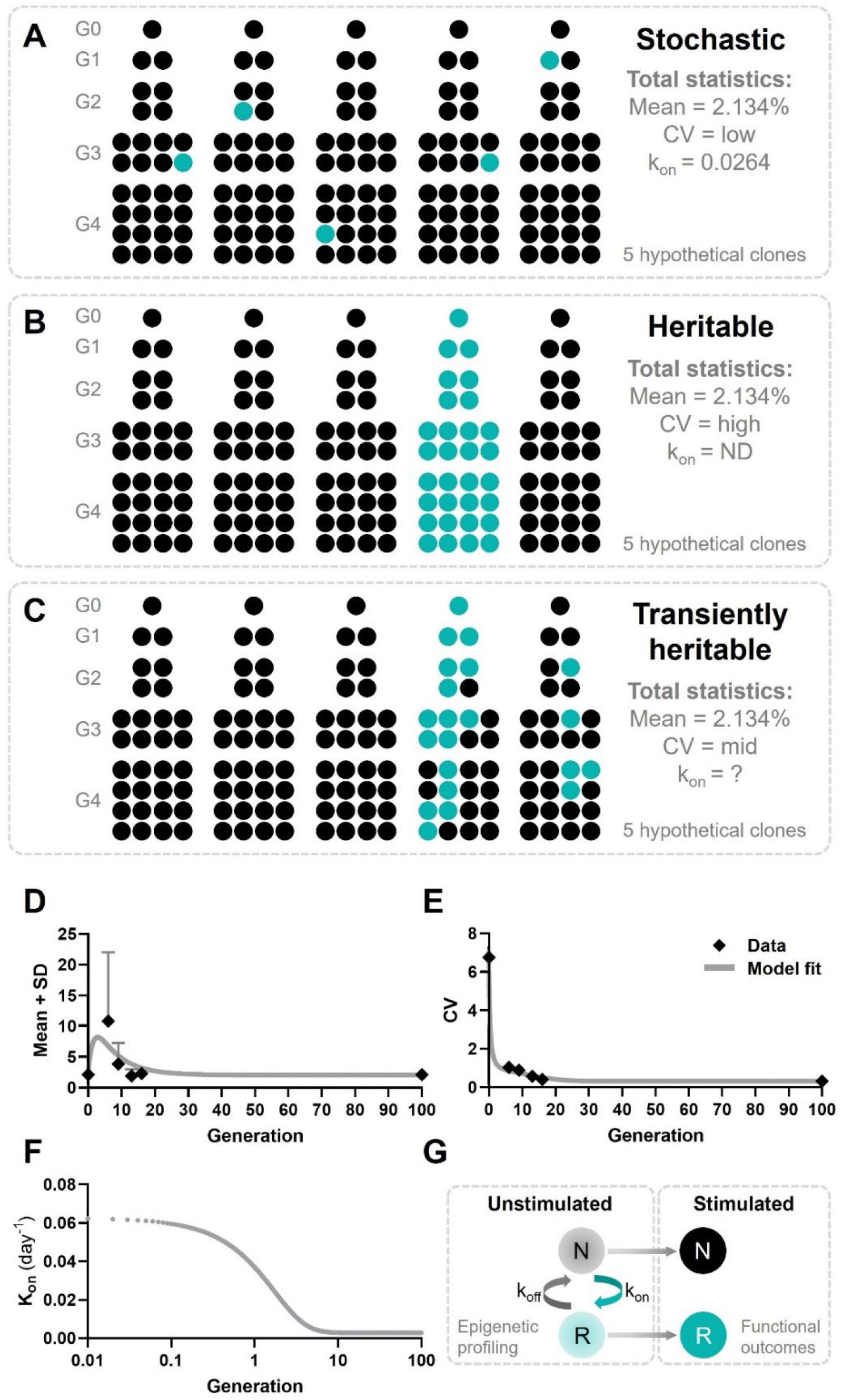
Modeling cellular decision-making during early IFN-I responses. **A** Hypothetical outcomes of responding cells upon cloning, assuming cellular decision making during early IFN-I responses is a stochastic process. Responders will appear randomly across clones, resulting in a total mean of 2.134%, a low coefficient of variation (CV), and a low k_on_ **B** Hypothetical outcomes of responding cells upon cloning, assuming cellular decision making during early IFN-I responses is a heritable process. Responders will appear only from the lineage that started with a responding cell, resulting in all offspring becoming responders. The total mean will still be 2.134%, though the CV will be high and the k_on_ will be zero, as no cell switch fate. **C** Hypothetical outcomes of responding cells upon cloning, assuming cellular decision making during early IFN-I responses is a transiently heritable process. Responders are more likely to appear in lineages originating from a responding cells, but can also appear in lineages that started with a non-responding cell. Besides, responding cells can also disappear from responding lineages. This results in a total mean of responding cells that higher than 2.134%, with a high CV, and a variable k_on_. **D** Mean plus SD of experimental outcomes of fluctuation assay with ODE model fitted. **E** CVs of fluctuation assay with ODE model fitted. **F** Time-dependent k_on_ values used to fit the ODE model. **G** Schematic on cell fate switching, enabling transiently heritable functional outcomes upon stimulation. Data information: in (D, E), data are presented as mean ± SD.

While a purely stochastic cellular decision fate and a strictly heritable cellular decision fate cover two extremes, the phenomenon of transiently heritable cell fates is characterized by a type of heritability that falls between those two ends of the spectrum (Lu et al., 2021; Shaffer et al., 2020). In fact, transiently heritable cell fates cover an intermediate timescale, in which cellular states may persist for several cellular divisions but are ultimately transient, and thus not indefinitely heritable. Still, this phenomenon can clearly be distinguished from the rather short-lived fluctuations referred to as noise (Shaffer et al., 2020). As a transiently heritable phenomenon allows responders to appear from non-responding parental cells, the mean across clones will still equal to 2.134%, while the CV will be relatively high too, but not as high as compared to a strictly heritable fate (Fig. 6c). The kon will be based on the probability of the reintroduction of responding cells, which can be variable, but should per definition be slower than the rate of cell division.

Surprisingly, the data obtained from clones of generation 6 resulted in a mean higher than 2.134% (Fig. 6d), and a relatively high CV (Fig. 6e). Accordingly, these results not only argue that the cellular decision to become a first responder is transiently heritable, but also indicate another underlying mechanism is causing this increased mean. From generation 13 onwards, both the mean and the CV start to meet the data obtained from the regular cultures again, which are similar to the theoretical outcomes of a stochastic process. One way to explain this clear difference, from a modeling point of view, is that that during the early generations the kon is much higher than during the later generations, reflected by the mean of 10.81% at generation 6. In other words, kon seems to be variable over the generations. As cell density is the only difference between generation numbers that the cells can possibly notice, we therefore assumed the kon to be density-dependent. Therefore, we fitted an ODE model, that incorporates this density-dependent kon variable (Fig. 6d, e). Details on the ODE model are provided in the Materials and Methods section. From generation 13 onwards, the overall population-wide response dynamics can be considered stochastic, with transiently heritable cell fates dictating responding lineages for only a few generations.

Together, we further confirm our hypothesis that cellular decision-making is defined by epigenetic profiling, which is transiently heritable, with the kon being density-dependent. Therefore, already in an unstimulated state, cells seem to switch between the two fates, which are reflected into functional outcomes upon stimulation (Fig. 6g).

### Quorum sensing drives cellular decision-making during early IFN-I responses

Following the observation of a varying kon, and our assumption that the kon is density-dependent, we next performed additional experiments aimed at elucidating the role cell density in dictating responsiveness. In practice, ranging from clones of generation 6 towards generation 13, the cell density (absolute cell count per area/volume) increases accordingly. This implies that at a lower cell density, corresponding with low generation numbers, cells tend to be programmed to become more responsive, meaning that percentages of responding cells becomes higher, and therefore the kon is higher.

The phenomenon of different cellular behaviors upon differences in cell density is in agreement with the concept of (immune) quorum sensing, which describes the ability of (immune) cells to perceive the density of their own population and adjust their behavior accordingly (Antonioli et al., 2019). Subsequent alterations in responsiveness are thought to be coordinated via epigenetic regulations. As we previously proved a role for epigenetics driving first responders, we wondered whether we could prove the effects of quorum sensing in cellular decision-making during early IFN-I responses. Therefore, we hypothesized that cellular decision-making is defined by epigenetic profiling, which switches over time between a responding and nonresponding state, even before stimulation, and is subject to the phenomenon of quorum sensing.

To test this final part of our hypothesis, we generated clones of generation 6 in low and high densities on coverslips as described before (Fig. 7a). We hypothesized that clones at low seeding densities display more fluctuations in the percentage of responders compared to high seeding densities, based on the results obtained in the fluctuation assay. Low seeding densities were obtained by seeding 250 cells per 24-well and verified upon visual inspection, meaning that these clusters of cells did not exceed the expected cell count of single clones (2^6/7^=64/128cells, depending on their grow speed), and were clearly separated from other clusters of cells, with over a 140 μm distance between the center points of the clones (Fig. 7b, c, Supplementary Fig. 7a). High seeding densities were obtained by seeding 1000 cells per 24-well, which resulted in merged groups of clones, thereby evidently exceeding the expected cell counts per cluster (Fig. 7d, e). In practice, clones seeded at high sending densities occasionally led to single clones, as observed upon low cell seeding (Supplementary Fig. 7b, c). For these instances, these clusters were considered as a single clones.

**Figure 7.**
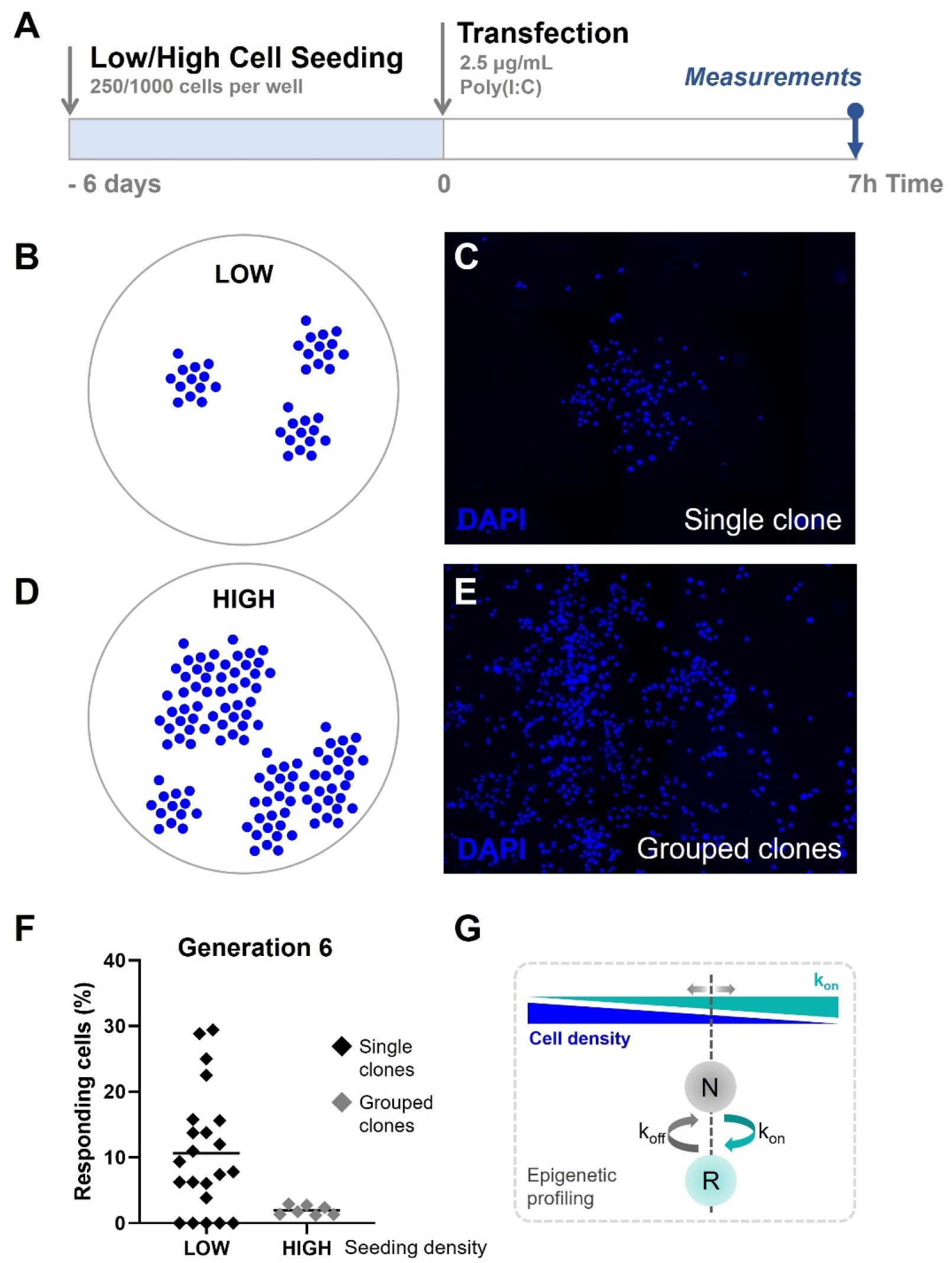
Quorum sensing drives cellular decision-making during early IFN-I responses. **A** Experimental design of quorum sensing experiments with NIH3T3: IRF7-CFP cells. Cells were either seeded at low numbers or high numbers (250 versus 1000 cells per 24-well) 6 days prior to the start of the experiment. Next, cells were transfected with 2.5 μg/mL Poly(I:C) and assessed for nuclear translocation of IRF7 after 7 hours. **B** Schematic representation of single clones of generation 6 on coverslips, seeded at low cell densities. **C** Microscopy image of DAPI channel, visualizing the nuclei of cells, displaying clear clustering of single clones of generation 6. **D** Schematic representation of grouped clones of generation 6 on coverslips, seeded at high cell densities. **E** Microscopy image of DAPI channel, visualizing the nuclei of cells, displaying grouped clusters of cells, consisting of numerous clones of generation 6. **F** Scatter plots on percentages of responding cell of clones of generation 6 seeded in low densities (n = 22 clones), and grouped clones seeded at high densities (n = 7). **G** Schematic on cell fate switching, influenced by cell density.

The results confirmed that single clones of generation 6 displayed high fluctuation, which closely matched with the data obtained earlier (average of 10.67 compared to 10.81; CV of 0.87 compared to 1.04, respectively) (Fig. 7f). Interestingly, the averages of merged clones of generation 6 displayed percentages of responding cells which closely matched with the numbers obtained from regular cultures (1.96% compared to 2.13, respectively).

To conclude, we confirm that cellular decision-making during early IFN-I responses is likely affected by the effects of quorum sensing. In other words, cell seem to be aware of their density, and adjust their epigenetic profiling to allow their secretory behaviors accordingly (Fig. 7g).

## Discussion

Here, we assessed the role of host-intrinsic factors dictating early IFN-I response dynamics. We observed that the cellular decision to become a first responder can be considered as a fate, rather than a coincidence driven by stochastic factors. Besides, this fate seems transiently heritable, regulated by epigenetic profiling and subject to the effects of quorum sensing. Because only a fraction of the cells become a first responder, quorum licensing might be a more suitable word of choice, as typically quorum sensing refers to a digital outcome in which either all cells or none at all respond (Muldoon et al., 2020).

Cells are faced by many decisions in response to external stimuli, reflected by a massive degree of cellular heterogeneity. By sharing information, a population of cells can make more effective decisions compared to each individual cell alone (Perkins and Swain, 2009). The ability of a fraction of first responders to drive population-wide IFN-I dynamics via paracrine signaling may be an efficient and robust strategy for quorum sensing, which allows tight regulation, but at the same time allows for flexibility and adjustability (Shalek et al., 2014). At the same time, this immune strategy is prone for mistakes. In autoimmune diseases like systemic lupus erythematosus (SLE), excessive IFNβ production potentiates auto-reactive dendritic cell activation (Hall and Rosen, 2010; Muskardin and Niewold, 2018). In contrast, excessively stringent thresholds may limit rapid responses to viral infection, as observed during severe acute respiratory syndrome coronavirus 2 (SARS-CoV-2) infection (Park and Iwasaki, 2020).

The phenomenon of a small fraction of first responders, responsible for the rapid and robust production of IFN-Is, has been observed across species and cell types. Therefore, we consider our utilized murine cell model as a good immune-cell or generic tissue-cell alternative for characterizing the fundamentals of cellular decision-making. Future studies have to translate the fundamental findings obtained in this study towards *in vivo* situations, and potentially, towards clinical applications. Although the translation of single-cell work might seem challenging, because of the seemingly unnatural situations mimicked with single-cell work, we believe that the fundamentals of cellular decision-making are similar across numbers, scales, and systems. Accordingly, studies in small intestinal organoids report similar bimodal IFN responsiveness (Bhushal et al., 2017). Likewise, transcription factor Nuclear Factor Kappa B (NF-κB) translocation follow similar all-or-nothing (i.e. digital) response dynamics, are prone to epigenetic licensing, and corresponding responding fates can be considered as transiently heritable (Clark et al., 2021).

Previously, the overall consensus on how first responders were thought to be regulated was by stochastic regulation, or in other words randomly (Wimmers et al., 2018). This rare fraction of cells has been described as indistinguishable from the rest, except in their expression of core antiviral gene expression programs (Shaffer et al., 2020; Shalek et al., 2014). In contrast, we currently hypothesize that first responders are predetermined, meaning that prior to stimulation individual cell fates can somehow be predicted. Of note, this is different from lineage fates, as cells are able to switch fates, both across lineages as well as during their cellular lives (Shaffer et al., 2020).

The additional mechanisms underlying the transiently heritable fate to become a first responder remain mysterious. Further work will be required to assess potential modules of gene regulation dictating these rare cellular decision-making processes, such as methylation or other regulatory mechanisms that operate on intermediate timescales (Meir et al., 2020). Regarding the phenomenon of quorum sensing, studies have reported that dendritic cell (DC) activation by poly(I:C) harbors a collective production of IFN-I, which drives DC activation at the population level in vivo (Bardou et al., 2021). Therefore, sustained IFN-I signaling in mediating full DC activation promotes collective behaviors, instead of cell-autonomous activity. In other words, the concentration of IFN-Is produced by a single cell has no biological effect, whereas the accumulation of IFN-Is produced by many cells drive a collective response. This makes us hypothesize that a low cell density, as occurs for clones of generation 6 in the fluctuation assay, increases the percentage of first-responder fates, thereby avoiding the risk of IFN-I levels that are too low to have any biological effect.

While transcriptional regulators have been the main focus of studying IFN-I dynamics, insights on additional types of regulation, such as epigenetic regulators and quorum sensing are shedding their light on an already complex IFN-I system. Although the presence of first responders mainly got characterized using microfluidic techniques, which seem far from representing the complex *in vivo* situation, studies have proven their existence and importance *in vivo* (Bauer et al., 2016; Zhang et al., 2020). Additionally, understanding the fundamentals of cellular decision-making during early IFN-I responses open compelling avenues for future development of novel IFN-I-targeted therapies. Especially considering the crucial role of well-orchestrated IFN-I response dynamics in clearing SARS-CoV-2 infection, while preventing harmful and ineffective cytokine storms (Park and Iwasaki, 2020), emphasizes the necessity of understanding the fundamentals of cellular decision-making. Together, the combination of single-cell technologies, mathematical modeling approaches, and the *in vivo* validation and translation continues to unravel the complexity of the IFN-I system in physiological contexts.

## Materials and Methods

### Cell culture and activation

Reporter murine fibroblastoid NIH 3T3 cells with stable expression of IRF7-CFP, STAT1-CFP and STAT2-YFP fusion proteins were cultured under standard tissue culture conditions in DMEM medium (Sigma) supplemented with 10% fetal calf serum, glutamine, penicillin, streptomycin, and selection antibiotic G418 or puromycin. Additional details on plasmids, BAC constructs, DNA transfections, and cloning are provided by literature (Rand et al., 2012). For experiments, cells were seeded on glass coverslips in 24-well plates, and activated using Lipofectamine2000 (Invitrogen) transfection reagent according to the manufacturer’s instructions. At all times, fluorescently labeled stimuli (rhodamine-labeled LMW Poly(I:C), InvivoGen) were used to assess transfection timing and efficiencies throughout the experiments. For additional transfection optimization, cells were measured with a flow cytometer (FACS Canto II, BD).

### Image and data analysis

Coverslips with cells were thoroughly washed (3x) with medium containing 10% FCS to loosen sticky liposomes from the glass and from the cell’s surfaces, to avoid false positivity upon assessing transfection efficiency. Next coverslips were fixed with 3% formaldehyde for 15 minutes at room temperature, washed, and stained with Hoechst 33343 to visualize nuclei. Next, coverslips were mounted on microscopy slides using Vectashield mounting media (Vector Laboratories), and imaged with a Nikon Eclipse Ti2 fluorescent microscope (Nikon). Image acquisition was performed by making multi-tile images at a magnification of 20x. Images were analyzed with ImageJ (National Institutes of Health) and a customized CellProfiler script (www.cellprofiler.org). Transfection efficiencies were determined based on the mean intensities provided by the CellProfiler script. The transfection threshold was based on the maximum intensity obtained from the untransfected cells. IRF7 translocation ratios were calculated using the following equation:

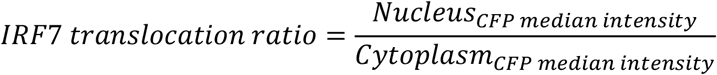

Images from which the percentage of translocated cells were drawn were at all times manually and visually checked, considering the relatively low percentages. Data visualization and statistical analysis were performed using the GraphPad Prism software (GraphPad).

### Fluctuation assay

Single cells were seeded into 96-wells plates using limited dilution in regular growth medium supplemented with 20% fetal calf serum and 20% condition medium obtained from regular cultures. Upon cell stretching, all wells were visually inspected to detect multiple seeded cells per well, and excluded from the experiments. For 6^th^ and 9^th^ generation clones, cells were seeded on glass coverslips in a concentration of 10 or 50 cells per well, respectively, and tracked over time to assure single-cell clones. Probability calculations were performed using the following equations:

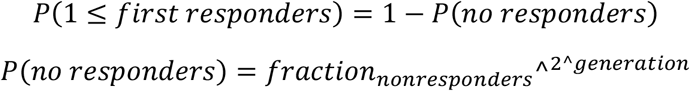

### Mathematical modeling

For this study we considered a simple model in which single cells can be in either one of two states: responder and nonresponder. Cells in the nonresponder state become responder with rate *k*_*on*_, and responders become nonresponder with rate *k*_*off*_. Based on the experimental data obtained from the fluctuation assay, we consider a time-varying *k*_*on*_ *(t)* that decreases with increasing colony size. We phenomenologically model this time-varying *k*_*on*_*(t)* as 

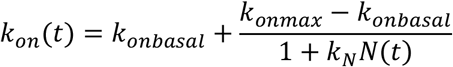

where *N(t)* =*e*^*rt*^ is the cell number over time and *k*_*N*_ is a positive constant. Note that as *t*→ ∞, *k*_*on*_*(t)* → *k*_*onbasal*_ and the fraction of responder cells approaches

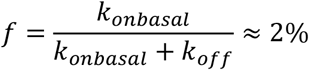

Based on the cell count at day 6 and 9, we estimate *γ* =0.73 day^−1^.

The average fraction of responders *x*(*t*) over time is given by the ordinary differential equation

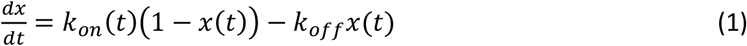

with an initial condition *x*(0) =2%. To predict colony-to-fluctuations in the fraction of responders, we assumed a Bernoulli initial condition of the starting single cell which is in the responsive state *x(*0) =1 with probability *f*, and in the unresponsive state *x(*0) =0 with probability 1- *f*. Let *x*(*t*)|_*x*(0)=1_ be the solution to the differential equation (1) with *x(*0) =1, and *x*(*t*)|_*x*(0)=0_ be the solution with *x*(0) =0. Then, the inter-colony coefficient of variation *CV* in the fraction of responsive cells is given by

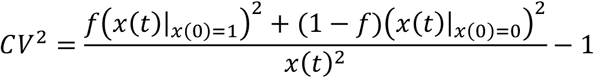

To account for the technical and biological noise, we further modify this equation to

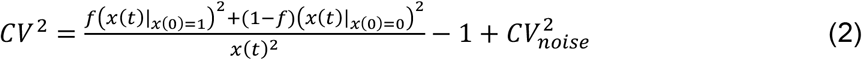

where *CV*_nois*e*_ ≈ 0.32, which is the fluctuation in the fraction of responders between independent samples from regular cultures. We fitted the solution of equation (1) and (2) to the fluctuation data to estimate parameters *k*_*onmax*_, *k*_*off*_ and parameter *k*_*N*_ corresponding to the above-defined function capturing the decreasing *k*_*on*_(*t*). For a given value of *k*_*off*_, *k*_*onbasal*_ is chosen so to ensure 2% of responders. Our fitting showed *k*_*off*_ =0.14 day^−1^ which corresponds to approximately 7 days in the responsive state before switching back to the unresponsive state. The time-varying *k*_*on*_ (*t*) obtained with this approach is shown in Fig. 6f.

## Acknowledgements

The authors would like to thank Ulfert Rand, Hansjörg Hauser, and Mario Köster for providing the reporter cells. Additionally, the authors would like to thank Nidhi Sinha and Bart M. Tiemeijer for the enthusiastic, insightful and lively discussions. This work was supported by the European Research Council (ERC) under the European Union’s Horizon 2020 research and innovation program (Grant agreement No. 802791). Finally, the authors would like to acknowledge the generous support by the Eindhoven University of Technology.

## Author contributions

LE, AS and JT designed the study. LE and VV performed experiments and analyzed the data. LE and AS performed the modeling. LE, AS and JT wrote the article. CB and JT supervised the research. JT acquired funding.

## Conflict of interest

The authors declare that no conflict of interest exist.

## Data availability

The raw data supporting the conclusions of this article are available on DataDryad.

## Supplementary Figures

**Supplementary Fig. 1.**
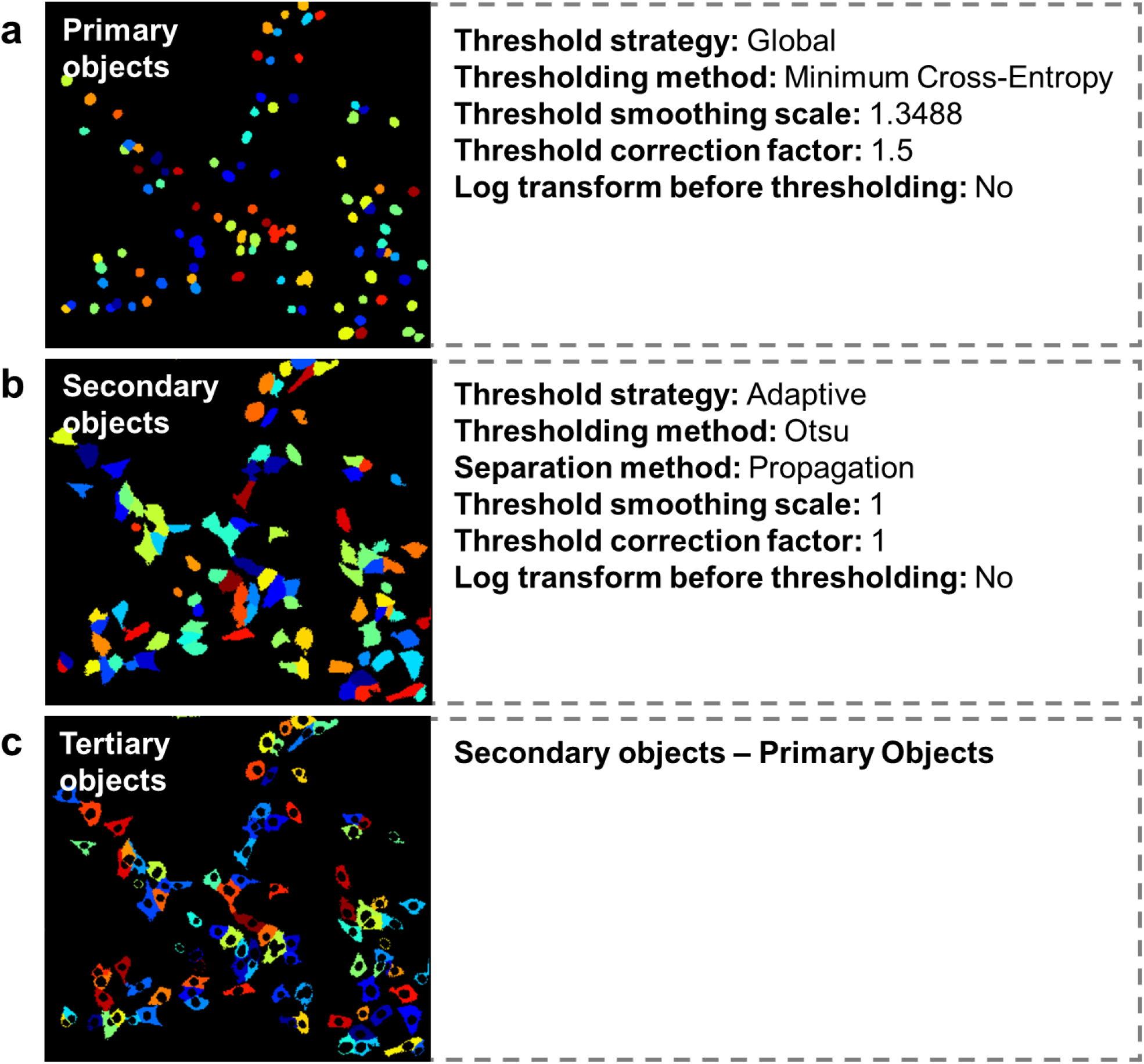
Details on automated script in CellProfiler software. **a-c** Identification of primary, secondary and tertiary objects.

**Supplementary Fig. 2.**
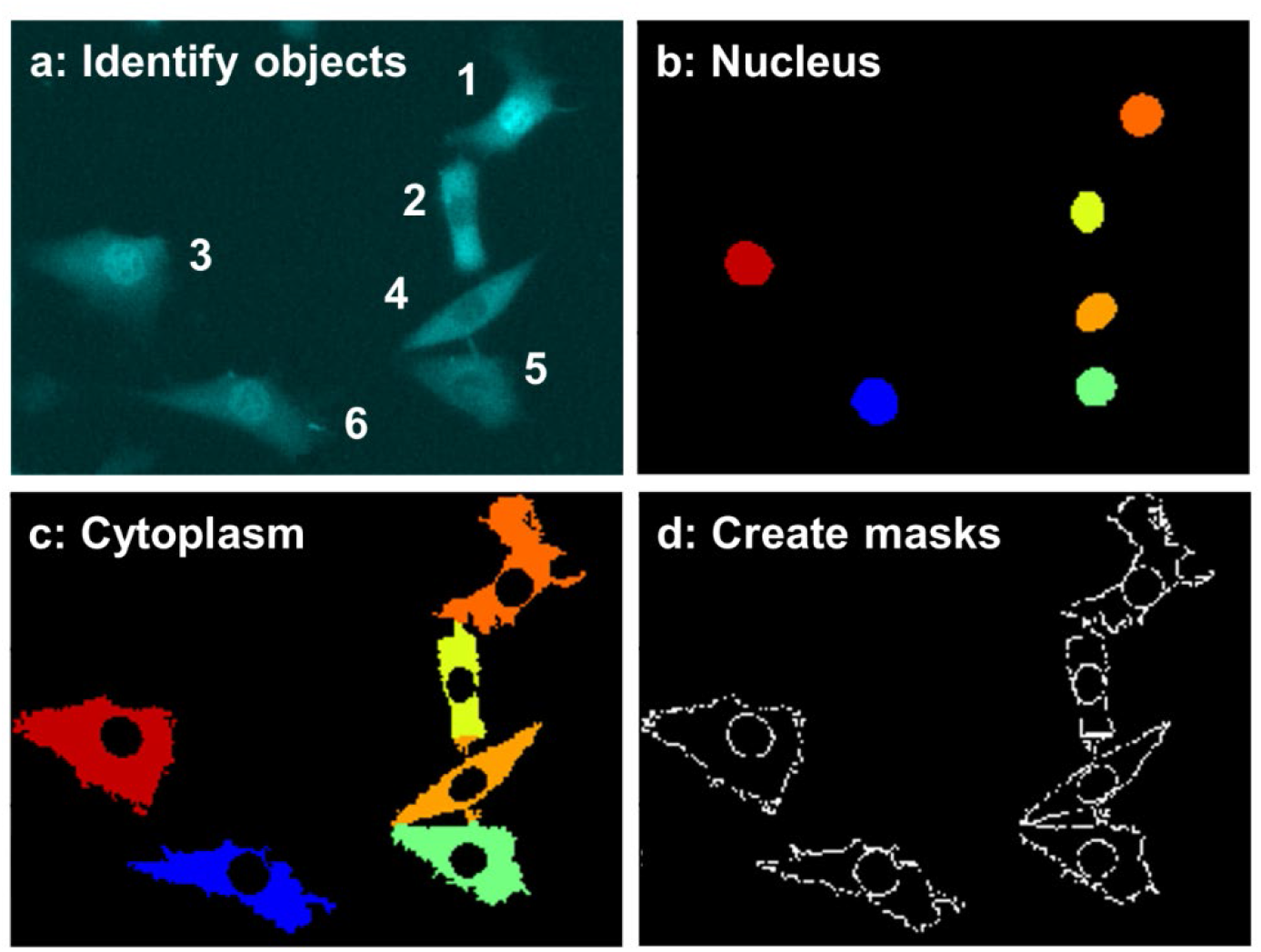
Detailed automated script analysis performance. **a** Identification and indexing of objects. **b** Detection of nuclei. **c** Detection of cytoplasm. **d** Creation of masks aligning the nuclei and cytoplasm.

**Supplementary Fig. 3.**
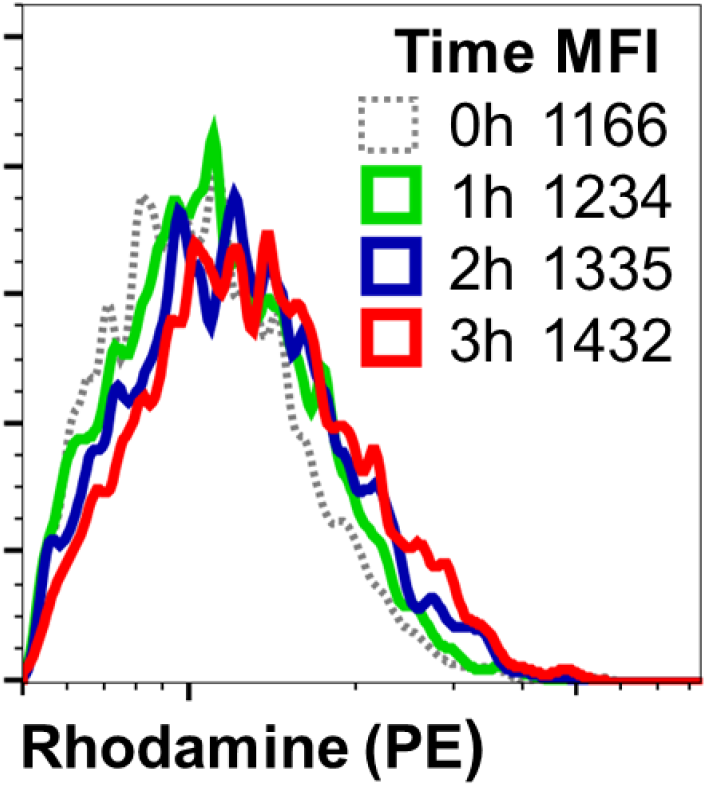
Flowcytometric analysis of transfection efficiency. Fibroblasts were transfected as described previously, incubated for up to 3 hours, during which for each hour the cells were trypsinized, thoroughly washed and measured. Depicted are the total fibroblast events, for unstimulated (0h), and for the first 3 hours after transfection, and their corresponding fluorescent mean fluorescent intensity (MFI) values for PE-Rhodamine.

**Supplementary Fig. 4.**
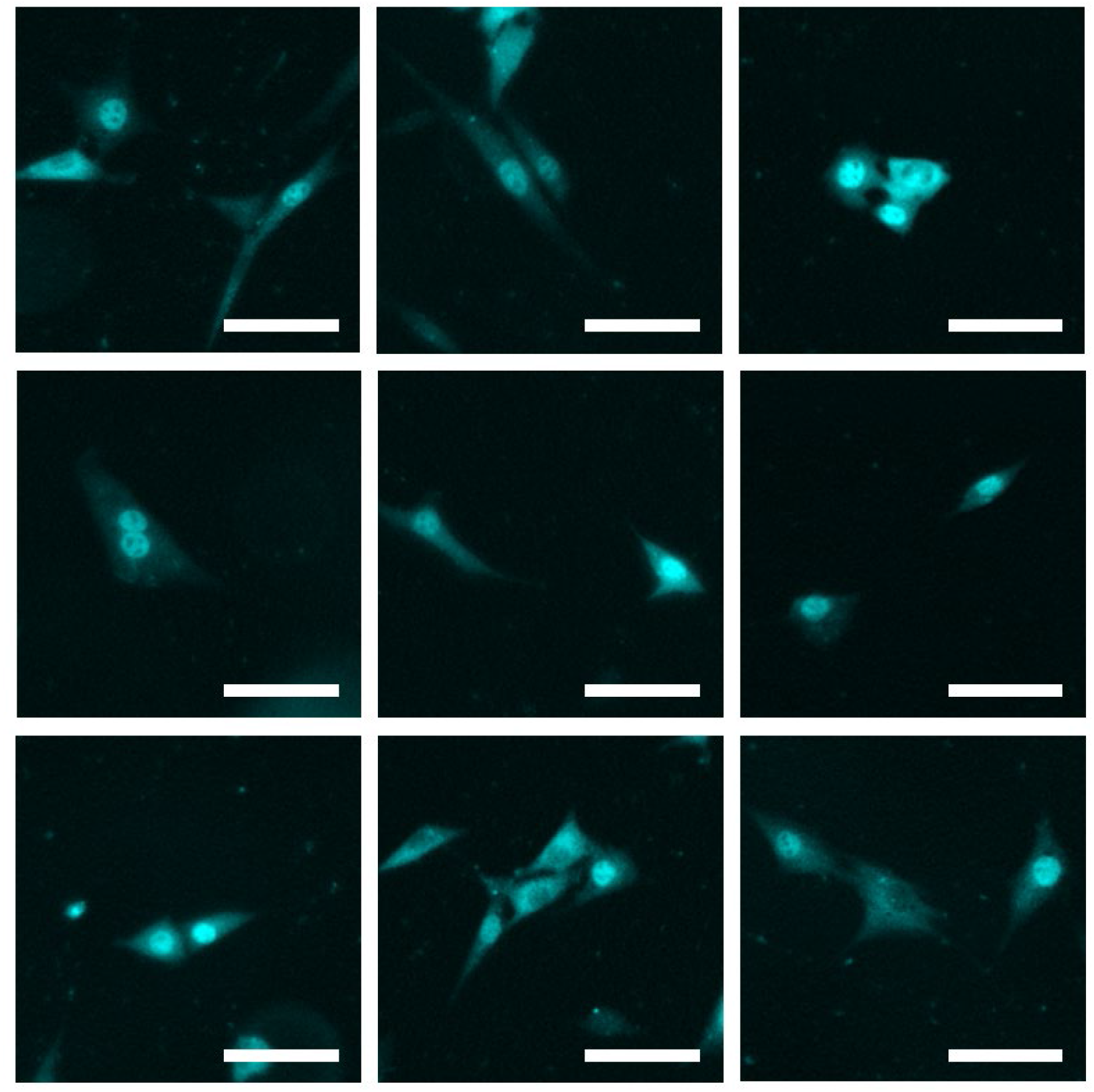
Microscopy images of numerous neighboring cells showing translocation. Several examples of neighboring cells showing translocation, transfected with 2.5 μg/mL Poly(I:C) for 7 hours, imaged and analyzed for IRF7 translocation. +20% Brightness and +20% contrast were applied for visualization purposes. Scale bar, 100 μm.

**Supplementary Fig. 5.**
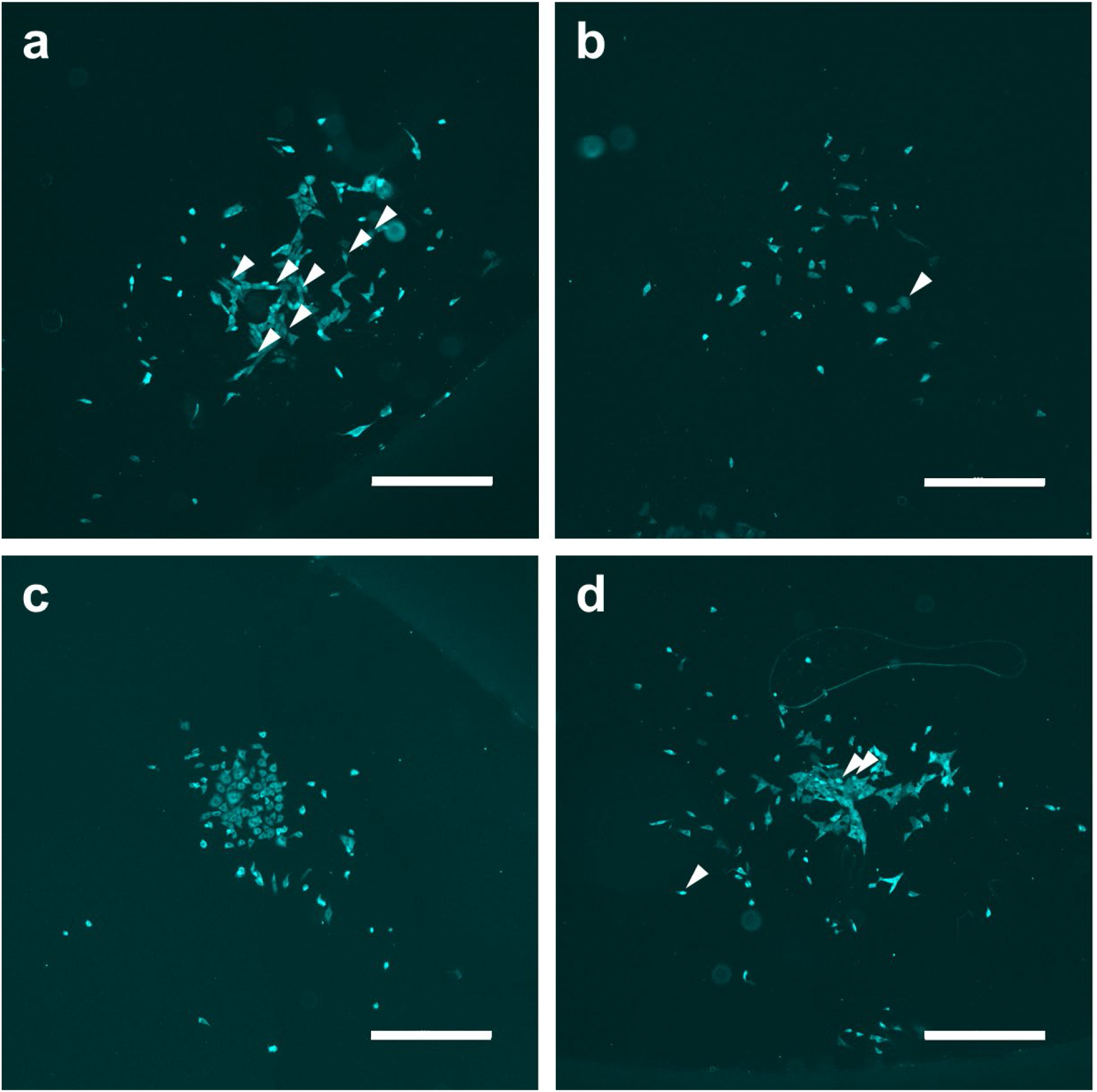
Microscopy images of clones of generation 6. **a-d** Several examples of clones of generation 6, transfected with 2.5 μg/mL Poly(I:C) for 7 hours, imaged and analyzed for IRF7 translocation, displaying numerous translocated cells, some of which are indicated with white arrows. +20% Brightness and +20% contrast were applied for visualization purposes. Scale bar, 500 μm.

**Supplementary Fig. 6.**
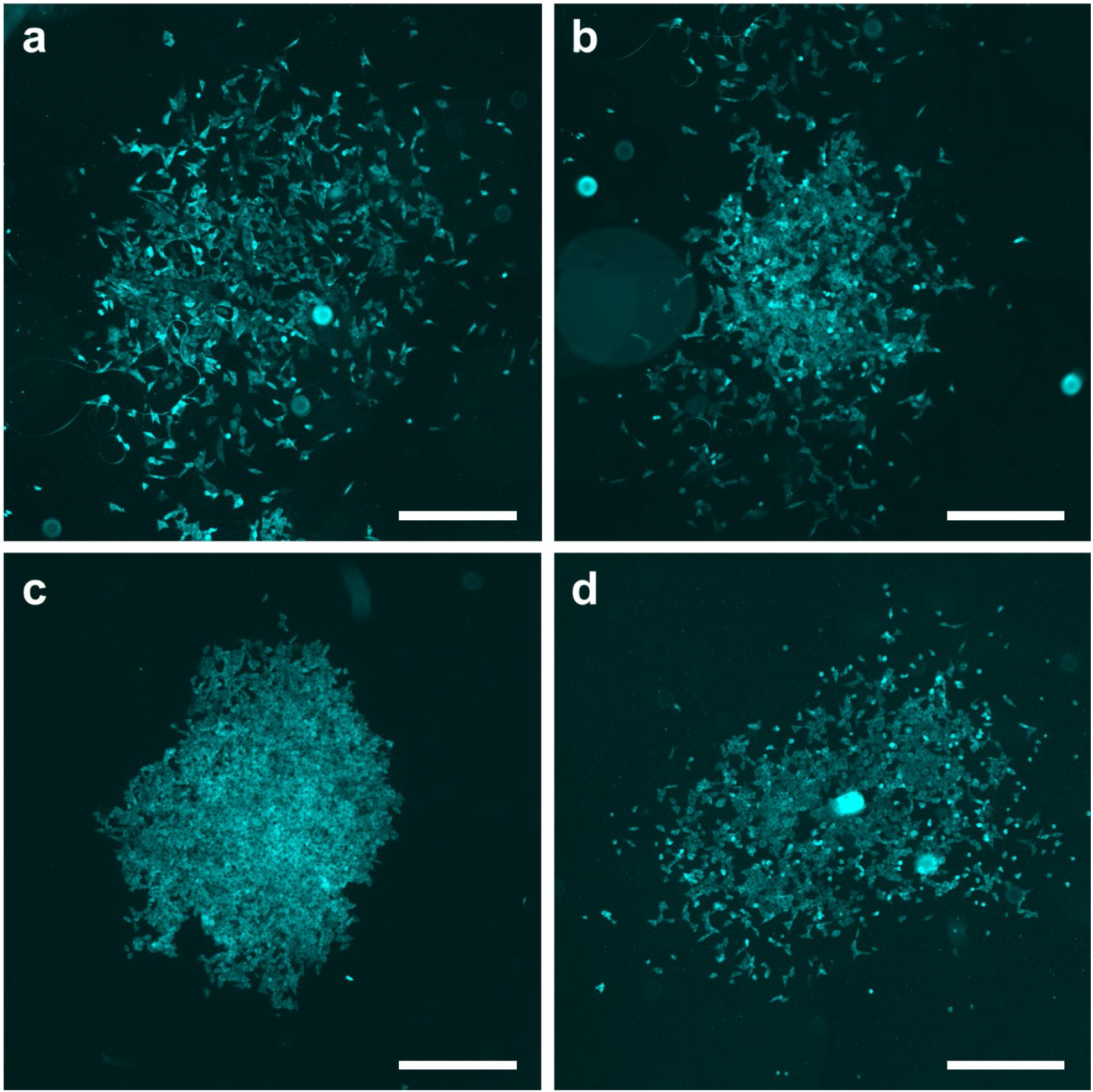
Microscopy images of clones of generation 9. **a-d** Several examples of clones of generation 9, transfected with 2.5 μg/mL Poly(I:C) for 7 hours, imaged and analyzed for IRF7 translocation. +20% Brightness and +20% contrast were applied for visualization purposes. Scale bar, 500 μm.

**Supplementary Fig. 7.**
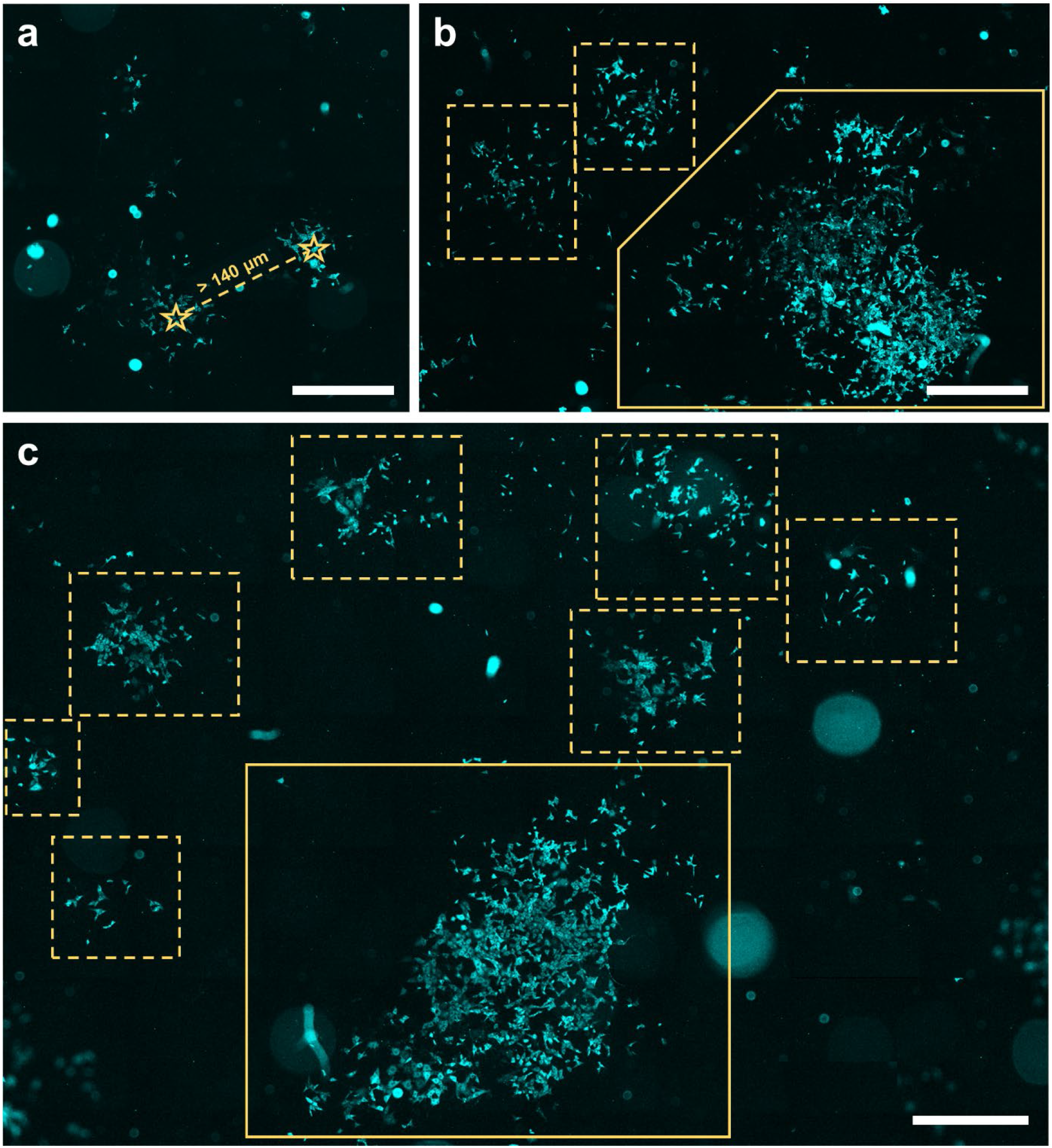
Microscopy images of cells of generation 6 seeded varying seeding densities. **a** Visualization of two clusters of cells that were considered two separate clones, with over 140 μm distance between the centers of the clusters, depicted by the star symbol. **b, c** Several examples of clones of generation 6, in dashed boxes, with in the same field a grouped clone, in solid box. +20% Brightness and +20% contrast were applied for visualization purposes. Scale bar, 1000 μm.

## Notes

### Competing Interest Statement

The authors have declared no competing interest.

